# vsgseq2: an updated pipeline for analysis of the diversity and abundance of population-wide *Trypanosoma brucei* VSG expression

**DOI:** 10.1101/2025.09.12.675796

**Authors:** Guy R Oldrieve, Stephen Larcombe, Marija Krasilnikova, Monica R Mugnier, Keith R Matthews

## Abstract

*Trypanosoma brucei* is an extracellular eukaryotic parasite that causes sleeping sickness in humans and Nagana, Surra and Dourine in livestock, game animals and horses. The parasite displays an extensive immune evasion mechanism, utilising the expression and ability to switch antigenically distinct variant surface glycoprotein (VSG) coats. VSG encoding genes account for ~10% of the *T. brucei* genome, and mosaic VSGs, assembled from distinct incomplete VSG gene copies, can be produced from this VSG library, generating an almost infinite VSG repertoire, which enables chronic infections. Each parasite expresses just one VSG at a time, but within a host, many VSGs can be expressed simultaneously. VSGSeq is an amplicon sequencing approach that enables surveillance of the population-wide diversity and abundance of expressed VSGs. vsgseq2 is an updated bioinformatics pipeline that enhances the reproducibility, accuracy, and efficiency of VSGseq analysis, utilising publicly available analytical tools.

**Plain Language Summary:** African trypanosomes, such as *Trypanosoma brucei*, are parasites that cause deadly diseases in humans and livestock. They survive in their host’s blood by constantly changing a protective coat of proteins, known as variant surface glycoproteins (VSGs). Switching VSG makes it very hard for the immune system to keep up, allowing infections to last for months or even years. At any one time, each parasite uses only one VSG, but across the whole population inside a host, many different VSGs are used simultaneously. To study how parasites change their coats, a method called VSGSeq was developed, which shows the genetic basis that makes the VSG coat. This research introduces vsgseq2, which provides an enhanced workflow for population-scale analysis of VSG expression, helping future research to understand how trypanosomes evade their host’s immune attack.

## Introduction

The single-celled parasite *Trypanosoma brucei* has evolved elaborate and divergent genome biology, most notably exemplified by its intricate mechanisms of immune evasion. As the parasite lives extracellularly, it is continuously exposed to its host’s immune system. To avoid immune clearance, each parasite expresses a single variant surface glycoprotein (VSG) type from a large genomic repertoire ^1,2^. The VSG proteins are densely packed on the cell surface, acting as an immunodominant coat recognised by the immune system. However, since some cells in the population switch expression to a new variant, the population can be sustained despite continual immune recognition and clearance. Although only ~400 complete VSG open reading frames have been annotated in the *T. brucei* genome, the parasite can generate novel VSGs that enable immune evasion in chronic infections through the assembly of mosaic genes from incomplete pseudogenes.

The study of VSG diversity and abundance has been assisted by the development of a protocol for surveying population-wide VSG transcripts, VSGSeq ^4^. VSGSeq takes cDNA and uses PCR primers specific for the conserved 14-mer in the VSG 3’-UTR ^5^ and a spliced leader sequence ^6^ to perform VSG-specific transcript amplification. The resulting products are then sequenced, typically using short-read technologies.

VSGseq has been instrumental in significant discoveries in the field of trypanosome biology and antigenic variation ^7–9^, but since its creation, the bioinformatic tools used in the VSGSeq analysis pipeline have evolved considerably or been discontinued. Here, we present an update and refinement to the VSGSeq pipeline: vsgseq2, which improves the accuracy, reproducibility and efficiency of VSGSeq analysis, whilst also helping to ease pipeline distribution and usability.

## Methods

The code used in the following analysis is available here: https://github.com/goldrieve/vsgseq2/WOR. The vsgseq2 pipeline can be accessed here: https://github.com/goldrieve/vsgseq2

### Benchmarking data

Control libraries generated in Mugnier et al (2015) from seven Lister427 clones, each expressing a different VSG (Tb427VSG-3, Tb427VSG-2, Tb427VSG-17, Tb427VSG-417, Tb427VSG-629, Tb427VSG-11 or Tb427VSG-9) ^4^ (SRR1740498-SRR1740509) were used to benchmark vsgseq2 against VSGseq. These lines had a selectable marker at the promoter of the active expression site and were grown in vitro in HMI-9 with antibiotic selection to minimise in situ switching. Parasites were counted with a haemocytometer and mixed to generate two control libraries. Control Library A consisted of cells expressing VSGs in the following percentages of the entire cell population in the library (Tb427VSG-3 = 90%, Tb427VSG-2 = 10%, Tb427VSG-17 = 1%, Tb427VSG-417 = 0.1%, Tb427VSG-629 = 0.01%, Tb427VSG-11 = 0.001%, Tb427VSG-9 = 0.0001%). Control library B consisted of cells expressing the seven VSGs in equal proportions. Both libraries were created using 1×10^6^ and 1×10^7^ cells, and each library had been sequenced in triplicate.

The percentage that each assembled transcript contributed to the whole population was calculated. Transcripts below 0.01% of the population were removed, and the total number of transcripts above this threshold was counted for each sample and VSG quantitation method.

### Longitudinal in vivo mouse data

VSGSeq data previously generated from longitudinal infections of four mice (SRR1740510-SRR1740545) ^4^ were also analysed to highlight the impact of the analysis pipeline (VSGseq or VSGseq2) on the interpretation of experimental VSG expression data.

### Data processing

Benchmarking and in vivo data were analysed using three methods (Fig. 1):

**Figure 1:**
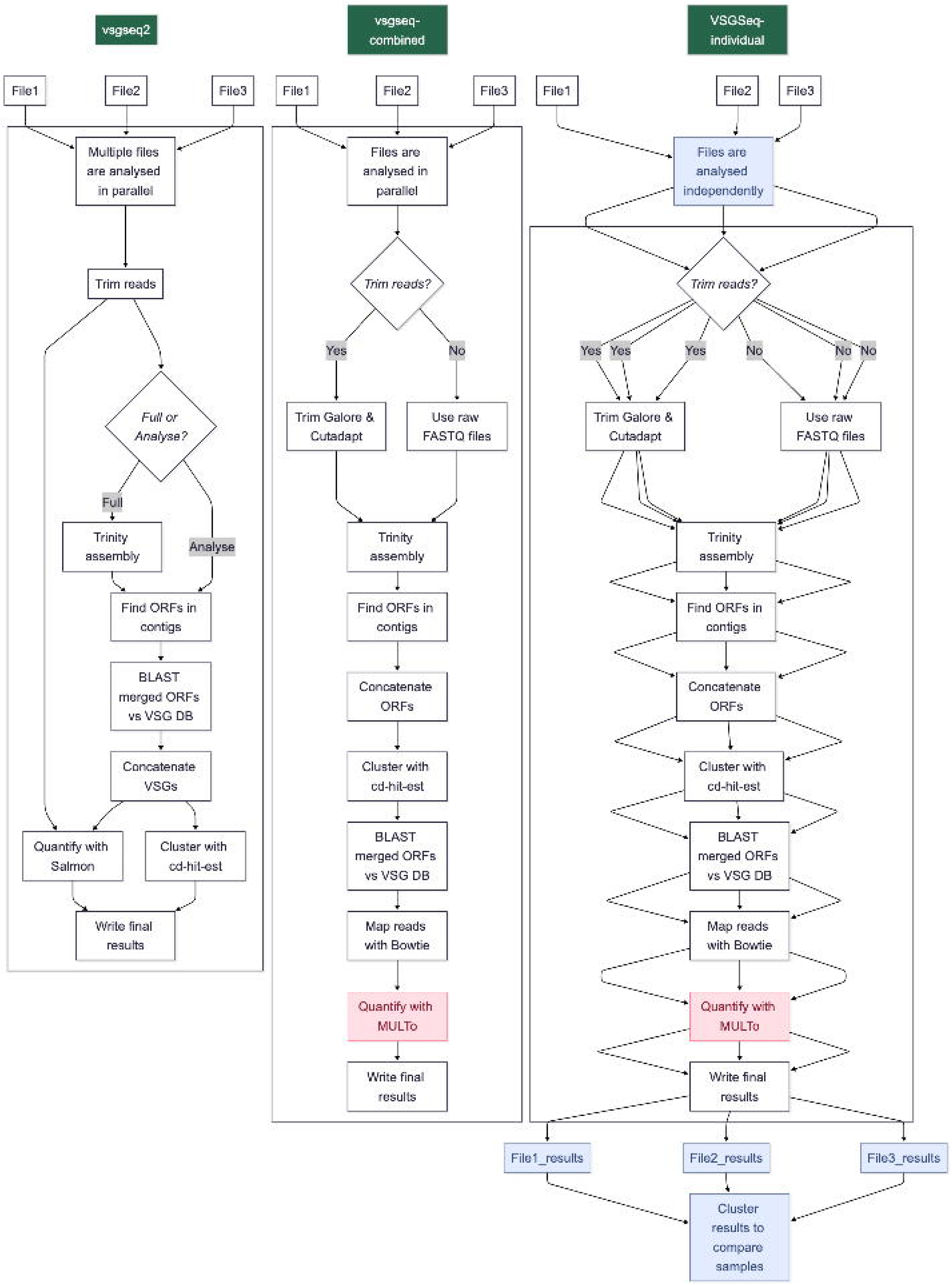
VSGSeq and vsgseq2 workflows follow similar analysis trajectories, ultimately turning VSG transcripts into assembled and quantified population-wide VSG expression surveys. In this study, benchmark and in vivo VSGSeq data were processed in three ways 1) VSGSeq was run over all samples together, allowing direct comparison between samples. 2) VSGSeq was run on each sample independently, and the results were then processed downstream of the pipeline to allow comparison between each sample. The code to run this method is not currently publicly available. 3) vsgseq2 was run over all samples in one execution. Here, the analysis steps performed during the three methods are visualised for three example samples. Large boxes represent analysis steps performed within a pipeline, and blue shaded boxes highlight those that need to be performed outside of a pipeline. Red shading highlights discontinued tools.

1. ‘VSGSeq-combined’: Currently, VSGSeq is recommended to be run by combining all samples from an experiment and executing the VSGSeq pipeline (https://github.com/mugnierlab/VSGSeqPipeline). This method combines all assembled VSGs from every sample and quantifies the expression of each assembled VSG for each sample.
2. ‘VSGSeq-individual’: To account for issues with the VSGseq pipeline when it was run in parallel across multiple samples, VSGSeq was run on each sample, and the data was then combined based on a cd-hit-est run on the individual sample runs.
3. ‘vsgseq2’: The pipeline is executed to process all samples from an experiment in combination, enabling direct comparisons between samples following completion of the pipeline.

To allow comparison between the three methods, the assembled VSGs were concatenated, and cd-hit-est was then used, with default vsgseq2 settings, to assign a cluster to each assembled VSG from the three methods, enabling us to visualise shared VSGs between each method.

### Pipeline distinctions between VSGSeq and vsgseq2

#### Pipeline structure

VSGSeq is written in Python 2, but the pipeline has not been maintained since the first release; the dependencies have undergone significant subsequent development and sequencing technologies have evolved. Specifically, VSGSeq uses Trinity-v2.8.5, MULTo has been discontinued and is no longer publicly available, and VSGSeq is restricted to the submission of single-end reads. Therefore, to update the dependencies and future-proof the pipeline, vsgseq2 was written and implemented using Nextflow ^10^, allowing for simplified pipeline distribution with its dependencies and modular design, assisting with future maintenance. In particular, the most computationally intensive process in VSGSeq analysis is the Trinity assembly. As vsgseq2 is modular, re-analysis of the analysis section of the pipeline, post-Trinity assembly, can be performed to remove the computationally intensive assembly. These can be initiated via the --mode flag (full or analyse).

#### Data preparation

Both VSGseq and VSGseq2 are executed by submitting a text file containing the sample ID and the location of the reads. The raw reads are then passed to Trim Galore for the removal of low-quality sequences and adapter sequences. Identical settings are used by both pipelines during the trimming step. In contrast to VSGSeq, vsgseq2 allows the submission of single-end or paired-end sequences at execution. The reads will be processed in parallel using identical tools, and specific settings are used for handling the different data types. This avoids the necessity in VSGSeq to concatenate paired-end sequencing reads together, which removes the benefits that paired-end information provides during Trinity assembly.

#### Transcript assembly and ORF prediction

Both vsgseq2 and VSGSeq use Trinity to assemble raw reads into transcripts. Similar default settings are used, but VSGSeq uses Trinity-v2.8.5, whereas vsgseq2 uses Trinity-v2.15.2.

In VSGSeq, ORFs are predicted using a custom Python script. Assembled transcripts are screened for start and stop codons, which are then extracted. However, if fully delineated ORFs are not found, the whole transcript is used as an ORF, such that partial ORFs can be produced. By default, vsgseq2 instead uses TransDecoder to predict and extract ORFs, with the –complete_orfs_only flag ensuring partial ORFs are removed. It is possible to include partial ORFs by executing vsgseq2 with the – partial_orf flag.

#### VSG identification

VSGSeq concatenates all ORFs into one file, performs cd-hit-est to remove duplicate sequences assembled in separate samples, and a blastn search is then performed for each of these sequences against a known ‘VSG’ database and a known ‘NotVSG’ database. The ‘NotVSG’ database consists of sequences manually annotated as false positives ^4^. In vsgseq2, identical cutoffs to filter VSGs from ORFs were retained, but the number of sequences in the ‘NotVSG’ database was increased, again through manual curation of false positive sequences.

In vsgseq2, ORFs are subject to BLAST, and the predicted VSGs are concatenated without performing cd-hit-est at this stage. Instead, there is direct progression to the next stage of the pipeline, where quantification of the expression of every predicted VSG occurs before performing cd-hit-est. In vsgseq2, cd-hit-est then picks a single VSG from a cluster of VSGs to represent all VSGs within that cluster; hereon referred to as the ‘champion’. Whereas in the VSGSeq pipeline, the champion is the longest sequence in a cluster of similar sequences, in vsgseq2, the VSG with the highest mean expression across all samples is selected to represent the ‘champion’ for that cluster. The result of these changes is that vsgseq2 chooses a champion sequence based on transcript abundance, rather than sequence length. Next, the TPM expression file produced by analysing all VSGs is summarised, and a new TPM file for the champion sequences is created. This is produced by summing the TPM for each VSG within a cluster, rather than re-running the quantification step.

#### VSG clustering and expression quantification

Analysis of benchmarking data containing sequencing reads from seven VSGs highlighted that VSGSeq is unable to assemble the correct number of VSGs when analysed in ‘VSGSeq-combined’ mode. Whilst this could be corrected by running ‘VSGSeq-individual’, this method still occasionally suffered from multiple versions of the same VSG being assembled and passed to the quantification stage, which led to the removal of multimapping transcripts. By analysing the benchmarking data and identifying cd-hit-est settings which would correctly combine independently assembled VSGs into the expected seven clusters, vsgseq2 was designed to correct the assembly of multiple versions of the same VSG. To achieve this, the cd-hit-est settings were altered in vsgseq2 to relax the cluster sequence identity, ultimately meaning more VSG sequences will be clustered and represented by a single sequence in the final VSGome (VSGSeq = cd-hit-est -i concatenated.fasta -o VSGome.fasta -d 0 -c 0.98 -n 8 -G 1 -g 1 -s 0.0 -aL 0.0 -M 50 -T 1 and vsgseq2 = cd-hit-est -i concatenated.fasta -o VSGome.fasta -d 0 -c 0.94 -n 8 -G 0 -g 1 -s 0.0 -aS 0.8 -M 50 -T 1).

VSGSeq-combined consistently used only a fraction of the generated sequencing reads due to a requirement for uniquely mapping reads in downstream quantification steps. VSGSeq uses bowtie and MULTo to quantify VSG expression, and default settings dictate that multi-mapping reads are removed from the analysis, presenting a significant problem when comparing complex VSG protein families. For example, VSGs often share large stretches of sequence identity, especially in mosaic VSGs, which can be formed from multiple donor VSGs. Removal of multi-mapping reads, in cases where multiple similar VSGs are present in a sample, can lead to a situation in which no reads uniquely map to a VSG, hindering the identification of VSGs even when they are present at high frequency within the population. This problem was especially apparent when VSGSeq was run in ‘VSGSeq-combined’ mode when similar VSGs were processed simultaneously in the VSGome quantification step. The result was that many dominant VSGs were under-represented in their apparent expression abundance, deviating from their underlying biological prevalence. In vsgseq2, Salmon is used to quantify VSG expression. Salmon assigns multi-mapping reads to the transcript with the most uniquely mapped reads, rather than discarding them ^11^. Although this will assign some reads to the wrong VSG, this trade-off is essential to gain a reproducible and accurate representation of population-wide VSG diversity and abundance, which is lost by the removal of multi-mapping reads. Furthermore, in many cases, the most biologically interesting expression changes occur among abundant VSGs. In vsgseq2, their accurate inclusion and quantification are weighted more highly than the potential loss of more minor transient VSGs from the analyses.

The issue of low sequence alignment rate can be resolved by running ‘VSGSeq-individual’. However, VSGSeq-individual requires post hoc merging of VSGs and quantification results. Furthermore, VSGSeq-individual is unable to handle multi-mapping reads from transcripts of very similar VSGs or variants of the same VSG.

Compared to VSGSeq-individual, vsgseq2 does not alter the ultimate biological signal extracted from a VSGSeq experiment, however, it does provide more consistent results across individual samples within an experiment and allows the scalable comparison of samples from any experiment directly following execution of the pipeline.

### Pipeline efficiency

Using the in vivo-derived VSGSeq data, the run time of each tool was calculated on a per-sample basis. The Green Algorithms calculator ^12^ was used to estimate the carbon footprint of these analyses. Default settings were used aside from the following: Type of cores = CPU, Number of cores = 1, Model = Xeon Gold 6248, Memory available (GB) = 20, Platform = Local server, Location = United Kingdom. The processing time consists of all steps post-Trinity assembly.

### Mosaic VSG data

To quantify the ability of each tool to process mosaic VSGs, sequences generated from a previous study, which identified expressed VSGs and putative donor sequences for the expressed mosaic VSG from a murine infection, were utilised ^13^. This data was based on cloning and Sanger sequencing the VSG from the expression site, therefore, the sequences were not derived from VSGSeq analysis. Marcello and Barry (2007) identified 21 expressed VSGs and generated a full sequence for 19. Of the 19 complete VSGs, 14 of the expressed VSGs could be accessed.

To test the pipelines, the 14 expressed VSGs with a putative donor sequence were each treated as a separate ‘sample’. Then, cd-hit-est was run on each ‘sample’ using the default cd-hit-est settings from VSGSeq or vsgseq2 and the number of sequences retained after cd-hit-est clustering was quantified and visualised as a heatmap.

The alignment of three ‘samples’ which were dealt with in different ways by the pipelines is presented as follows: 1) both vsgseq2 and VSGSeq cluster the donor or expressed sequence, 2) only vsgseq2 clusters a sequence and 3) both vsgseq2 and VSGSeq do not cluster sequences.

## Results

The benchmark and in vivo VSGSeq data ^4^ were reanalysed in three ways (refer to the methods section and Fig. 1 for more details):

1. VSGSeq-combined: Currently, VSGSeq is recommended to be run by combining all samples from an experiment and executing the VSGSeq pipeline. This method combines all assembled VSGs from every sample and quantifies the expression of each assembled VSG for each sample.
2. VSGSeq-individual: The outputs of VSGSeq run iteratively on individual samples are combined post-hoc based on cd-hit results using the normal pipeline parameters.
3. vsgseq2: Executed to process all samples from an experiment in combination, enabling direct comparisons between samples following completion of the pipeline.

### vsgseq2 improves the reproducibility and accuracy of population-wide VSG recapitulation

In an initial benchmarking experiment, Mugnier et al (2015) performed VSGSeq on seven clones, each clone expressing a unique VSG (Tb427VSG-3, Tb427VSG-2, Tb427VSG-17, Tb427VSG-417, Tb427VSG-629, Tb427VSG-11, Tb427VSG-9). The cells were either mixed in 10-fold dilutions (library A) or equal proportions (library B). Each library was generated using 1×10^6^ or 1×10^7^ cells and every iteration was sequenced in triplicate.

VSGSeq-combined, VSGSeq-individual and vsgseq2 were run on the benchmark data generated from these libraries and the number of VSGs above the limit of detection quantified, expecting to resolve seven VSGs. The original analysis of these samples utilised a pipeline similar to “VSGSeq-combined,” though with some differences in clustering parameters and software versions, which explains differences between these results and the previously published analyses of this dataset. VSGSeq-combined produced far more VSGs than both VSGSeq-individual and vsgseq2 for both library A and library B (Fig. 2a). This highlights that VSGSeq-combined does not perform well when run on multiple samples. Along with the overproduction of assembled VSGs, VSGSeq-combined cannot utilise the vast majority of the input sequencing data due to its requirement for uniquely mapping reads. Given the redundancy of the VSG repertoire and the parasite’s ability to generate mosaic variants, multi-mapping reads are an especially important consideration for VSG datasets, and this problem is amplified when mapping is performed using a single reference for multiple samples. vsgseq2 aligns significantly more reads than VSGSeq-individual (Fig. 2b).

**Figure 2.**
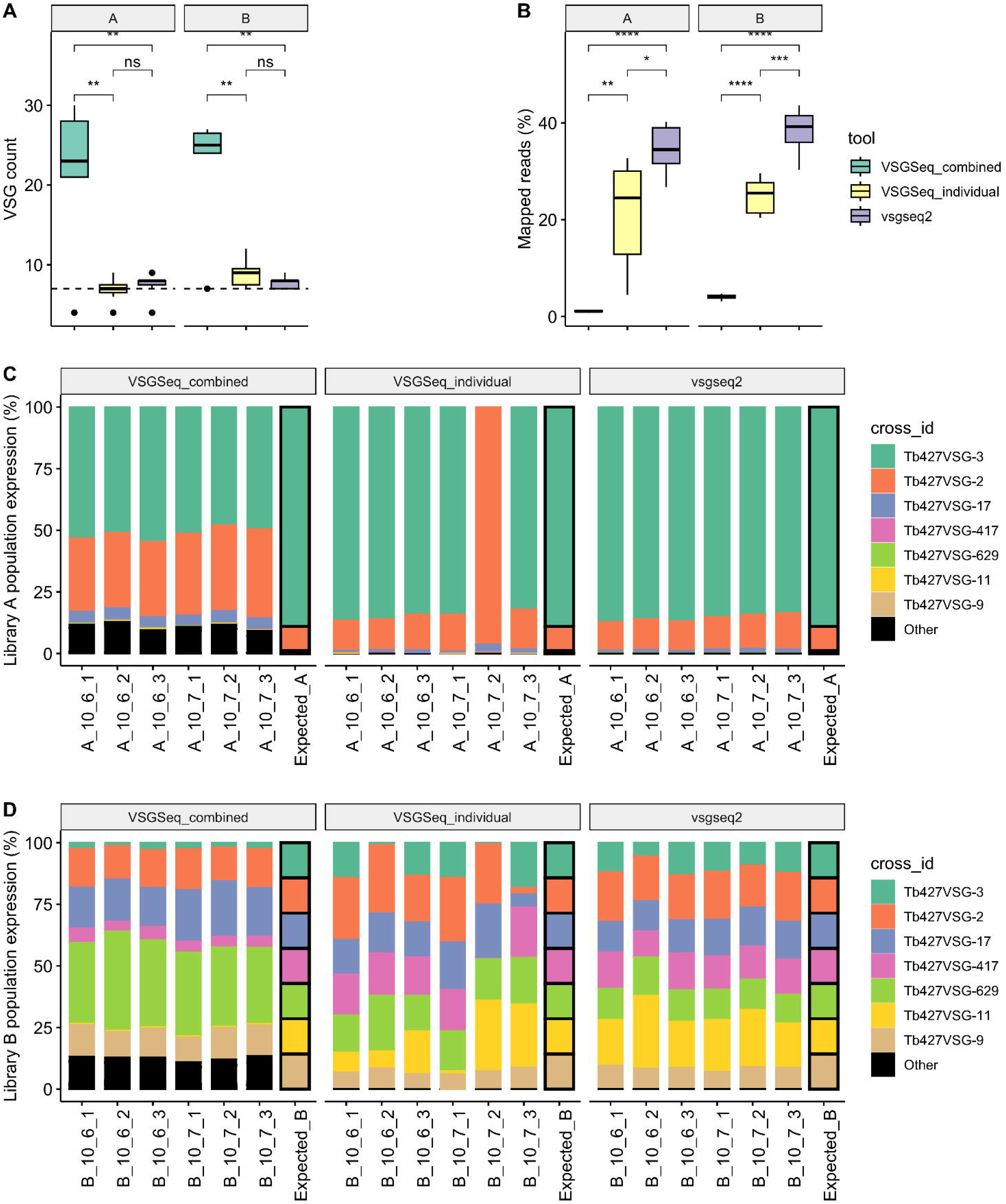
vsgseq2 accurately recapitulates benchmark VSGSeq data. **(a)** vsgseq2 detected the expected number of VSGs and **(b)** used significantly more of the input data compared to VSGSeq-combined and VSGSeq-individual. **(c-d)** This enabled a more accurate population-level quantification of VSG diversity and abundance. Each colour represents the percentage of each VSG cluster found in each sample for each analysis method.

The data were visualised to highlight how the overproduction of VSGs and the removal of multi-mapping reads altered the expression of the seven VSGs compared to the expected expression level. vsgseq2 recreated the expected proportions of the seven VSGs most accurately for both libraries A and B. For example, a considerable proportion of sequencing reads assembled by VSGSeq-combined were assigned to the ‘other’ (black) VSG bar (Fig. 2c), and VSGSeq-individual struggled to estimate the correct proportion of cells expressing Tb427VSG-3 in sample A_10_7_2 (Fig. 2d). In this specific sample run, multiple versions of Tb427VSG-3 were produced during Trinity assembly which led to removal of sequencing reads from the Tb427VSG-3 transcript from the analysis, as they mapped to two transcripts.

### Validation of vsgseq2 using in vivo data validates benchmarking results

Next, the VSGSeq data from longitudinal chronic infections of four mice were analysed ^4^. Blood samples were taken from day 6-7 until day ~30, except for mouse three, which survived until day 105. VSGSeq-combined overproduces VSGs (Fig. 3a), and vsgseq2 used significantly more reads than both VSGSeq-combined and VSGSeq-individual (Fig. 3b), highlighting VSGSeq’s requirement for uniquely mapping reads for quantification. Although vsgseq2 quantified a significantly higher numbers of reads, the mean mapping rate is still only 37%. Whilst the VSGSeq protocol enriches for VSG specific material, it is impossible to remove all host material, which leads to the low overall mapping rate. Due to the removal of multi-mapping reads, VSGSeq-combined often distorted the population-wide expression profile compared to vsgseq2 and VSGSeq-individual (Fig. 3c) and, in some cases, could not quantify expression of the dominant VSG due to an insufficient number of uniquely mapping reads, for example, M1-d24 (Fig. 3c).

**Figure 3:**
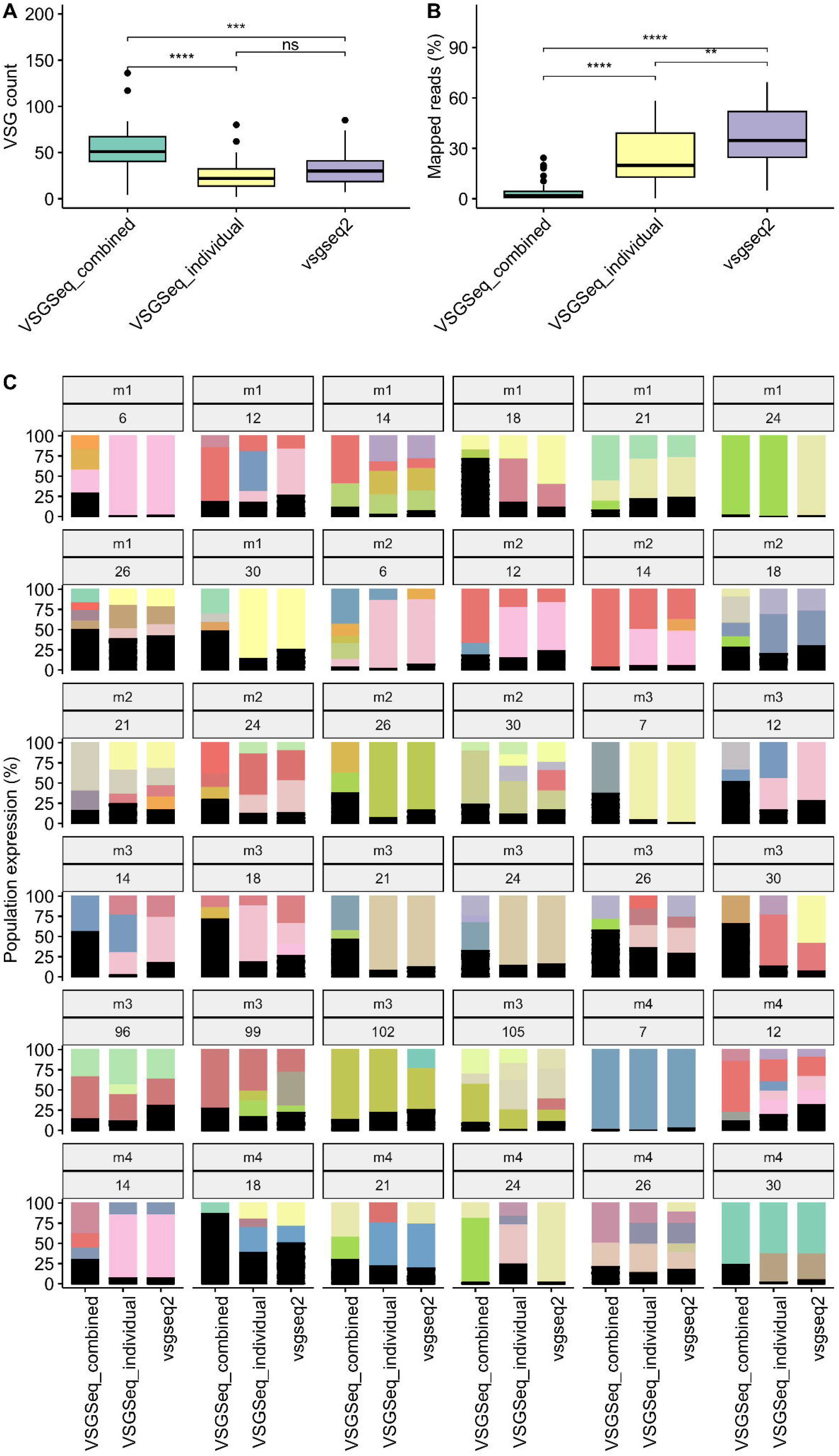
Analysis method impacts recapitulation of VSG diversity and abundance of in vivo infection experiments. **(a)** VSGSeq-combined produces significantly more VSGs than vsgseq2, and **(b)** VSGSeq-combined and VSGSeq-individual use significantly fewer reads to quantify the expression of these VSGs. **(c)** This causes variation in the predicted dominant VSG when the same data is analysed by VSGSeq-combined, VSGSeq-individual or vsgseq2. Each panel represents a VSGSeq sample taken from a mouse (mouse 1-4) across a longitudinal infection (day ~7 to 105). Each colour represents the percentage of each VSG cluster found in each sample for each analysis method.

During the pipeline update, the analysis efficiency was also improved. Ultimately, this led to a significant reduction in the runtime and carbon production for analysis of in vivo VSGSeq data (Fig. 4).

**Figure 4:**
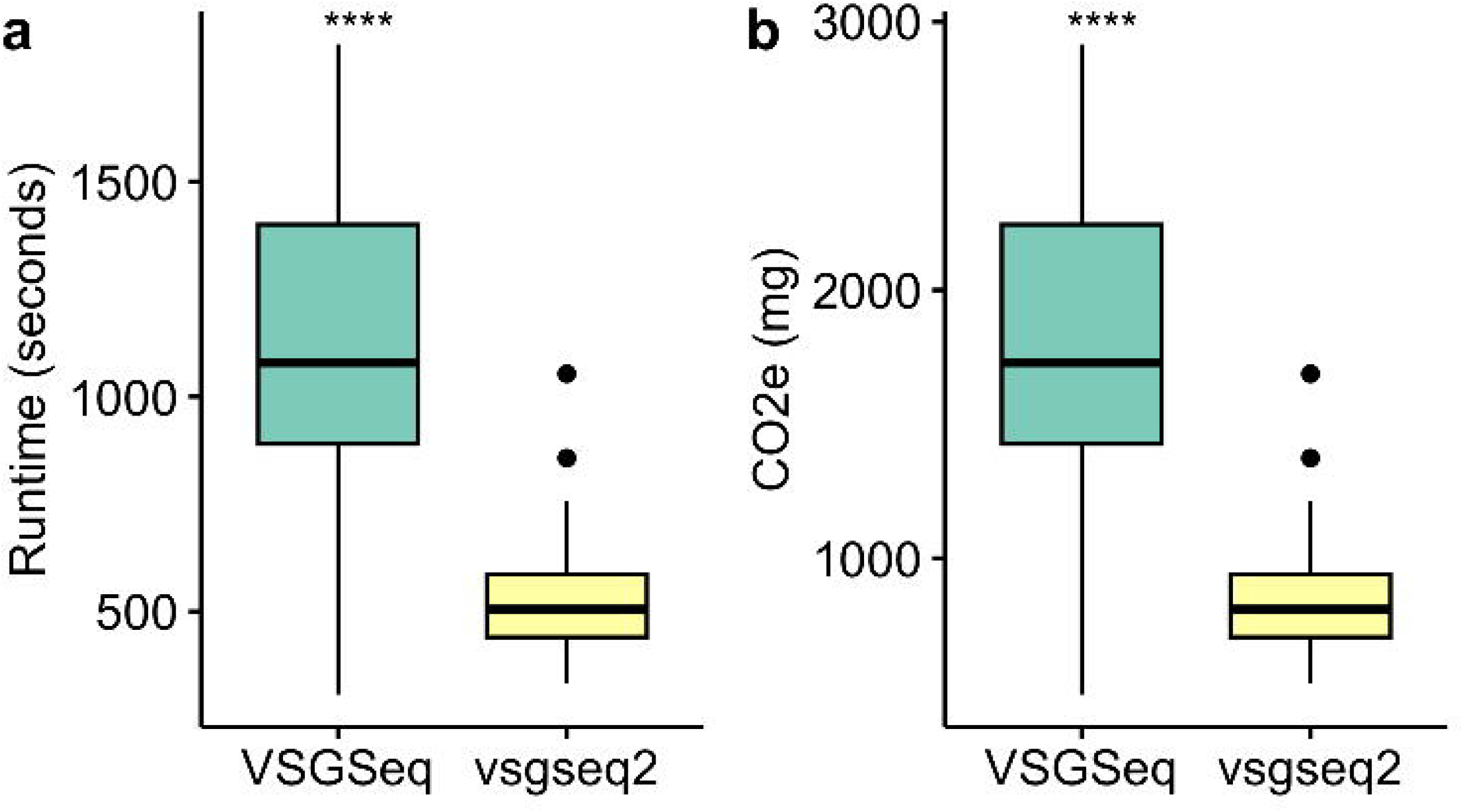
vsgseq2 is significantly more efficient than VSGSeq. The efficiency improvement enables a significant reduction in (a) runtime and (b) emissions produced when analysing the same VSGSeq data.

### VSGSeq and vsgseq2 both capture mosaic VSGs, vsgseq2 is more conservative

The cd-hit-est clustering settings used in vsgseq2 were calibrated to ensure seven assembled VSGs were produced from known benchmarking data produced from parasites expressing seven VSGs (Fig. 2a). To achieve this, the percentage identity used to cluster VSGs was reduced from 98% to 94% whilst also performing local alignments of the sequences rather than global alignments.

To test the impact of this change on the pipeline’s ability to capture mosaic VSG sequences, VSG sequences generated from a previous study that identified expressed VSGs and their putative donor sequences were downloaded. These sequences were not derived from VSGSeq analysis but were based on cloning the VSG from the expression site and Sanger sequencing of the resulting products. Marcello and Barry, (2007) analysed 11 mice, over a total of 28 days and identified 21 expressed VSGs; it was possible to generate a full sequence for 19. Of these 19 VSGs, we were able to find and download the putative donor sequences of 14 of the expressed VSGs.

Each of these 14 expressed and donor VSGs were treated as an independent ‘sample’, mimicking assembled VSGs created as part of the VSGSeq and vsgseq2 pipelines. Using cd-hit-est, these sequences were clustered from each of the ‘samples’ using the default cd-hit-est settings in VSGSeq and vsgseq2. In total, vsgseq2 performed identically to VSGSeq in 9 of the 14 expressed VSG ‘samples’ (Fig. 5a). Out of 14, four and eight ‘samples’ successfully progressed through the vsgseq2 and VSGSeq pipelines, respectively, without collapsing the donor sequences into a cluster with the expressed VSG.

**Figure 5:**
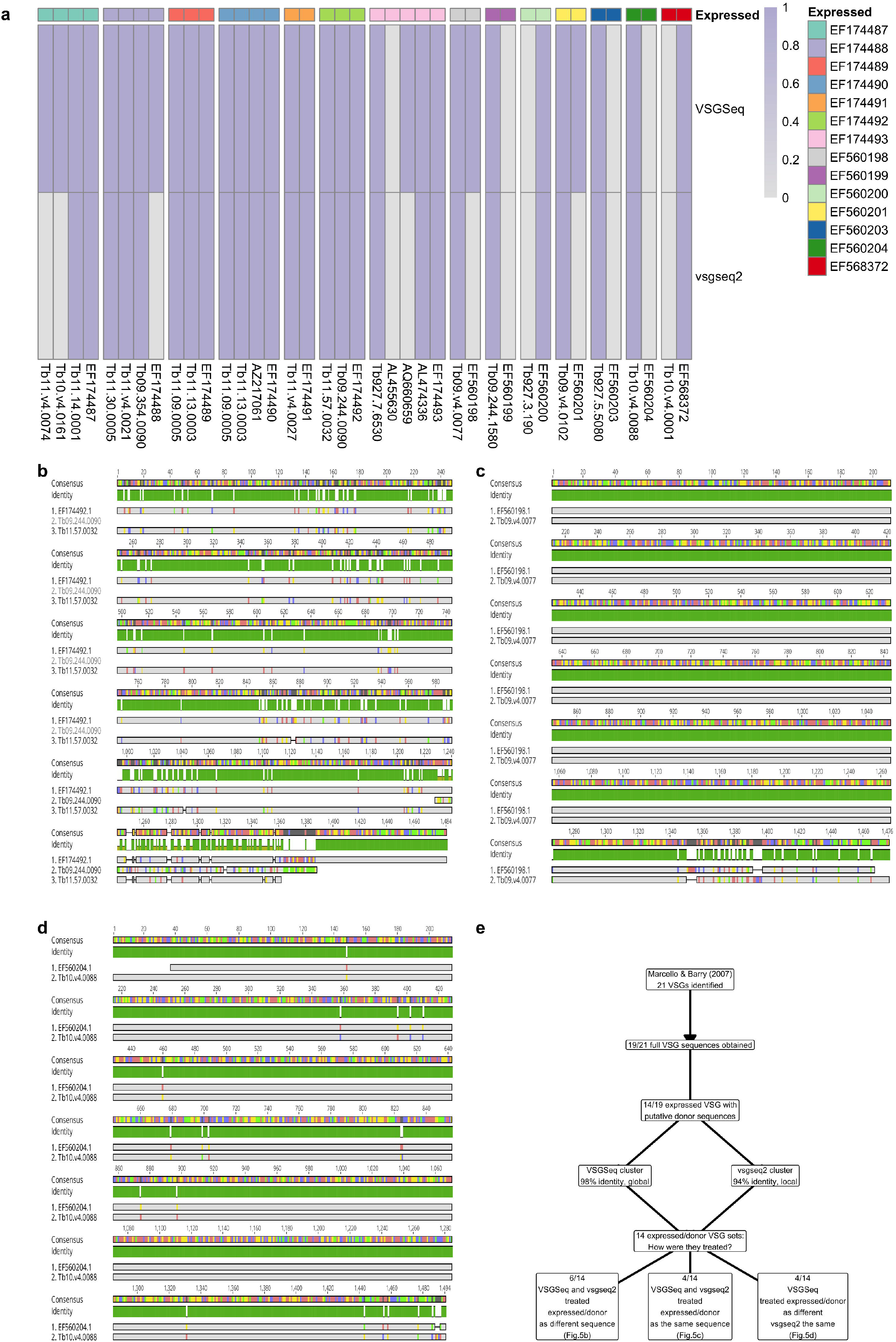
Both tools are capable of capturing mosaic and donor VSGs, but vsgseq2 is more conservative. **(a)** Presence and absence of expressed VSGs and their putative donor sequences during analysis with VSGSeq or vsgseq2. (**b-d**) Alignments of an expressed VSG and its putative donor for three cases: **(b)** both VSGSeq and vsgseq2 do not cluster the expressed or donor sequences and so the expression of both sequences would be quantified in the final ouput, **(c)** VSGSeq does not cluster the expressed and donor sequences, but vsgseq2 does, meaning both sequences would be quantified in VSGSeq but not in vsgseq2 and **(d)** both VSGSeq and vsgseq2 cluster the donor and expressed sequences, so neither pipeline would quantify both donor and expressed VSG.

The donor and expressed VSG sequences from three example ‘samples’ were aligned. Fig. 5b highlights a scenario where both VSGSeq and vsgseq2 did not cluster the expressed or donor sequences as the variation was high enough to classify them as separate VSGs. In Fig. 5c, VSGSeq does not cluster the expressed and donor sequences, but vsgseq2 does. In Fig. 5d, VSGSeq and vsgseq2 cluster the donor and expressed sequences, as there was not enough variation between the sequences to classify them as independent VSGs. Both pipelines were capable of analysing assembled donor and mosaic VSGs; vsgseq2 is more conservative based on control data benchmarking data (Fig. 2a).

## Discussion

VSGSeq facilitates population-wide surveys of VSG diversity and abundance, enabling an understanding of VSG expression and antigenic variation at the population scale. However, tool development has progressed substantially in recent years, and the VSGSeq pipeline required updates to maintain pace with these advances, and legacy components are no longer available or current. As an example, MULTo is no longer publicly available, and other essential tools, such as Trinity, have undergone seven years of further development. Mapping and quantification tools at the time of the development of the original VSGseq pipeline relied upon uniquely mapping reads for VSG quantification, with a correction for mappability using the tool MULTo. However, reproducibility was challenging when the number of samples being analysed in parallel was varied, and the assembly of similar VSGs and subsequent removal of multi-mapping reads could lead to the loss of expression data for VSGs, including dominant VSGs, from the results entirely. Whilst these difficulties can be mitigated by running VSGSeq on each sample individually, and then generating VSG clusters using cd-hit-est to allow comparison between these samples, vsgseq2 enables reproducible analysis of VSGSeq data whilst avoiding the requirement for downstream analysis outside of the VSGSeq pipeline.

Our analysis demonstrated that vsgseq2 accurately captures the diversity and abundance of expected VSGs from benchmarking data (Fig. 2). Whilst the outputs of VSGSeq-individual are closer to vsgseq2, vsgseq2, which utilizes the modern sequence alignment tool *Salmon*, can utilise the greatest percentage of sequencing data to recapitulate the population-wide expression, providing a more robust estimate of VSG diversity and abundance (Fig. 3). Furthermore, vsgseq2 analysis is significantly more efficient than VSGSeq, reducing the time taken to analyse data and reducing resource usage (Fig. 4).

To enable accurate numbers of assembled VSGs, the cd-hit-est settings were altered, which ultimately means more assembled VSGs are clustered. These parameters were based on the expected benchmark VSG assembly numbers (Fig. 2). Nonetheless, this may cause the removal of more similar VSGs, which could be true mosaics, rather than those that arise due to sequencing errors or misassemblies. Analysis of the ability of the pipeline to capture mosaic VSG sequences showed that both pipelines are capable of analysing assembled donor and mosaic VSGs, vsgseq2 is more conservative, based on control benchmarking data (Fig. 2a). Moreover, the tradeoff of being more conservative in regards to very similar VSGs is essential in identifying the dominant and often most important VSG in a sample. Whilst we have based the default settings in vsgseq2 on benchmark data, vsgseq2 is customisable, and the user can tune these settings for their analysis based on their experimental priorities.

In conclusion, based on its optimised analysis pipeline, ability to accurately identify VSG expression profiles in datasets of known VSG expression composition and the public and current accessibility of analytical modules, vsgseq2 provides an enhanced workflow for population-scale analysis of antigen gene expression in *T. brucei*.

## Data and software availability

### Source data

Raw sequencing data was accessed from NCBI SRA (SRR1740498 - SRR1740545).

### Underlying data

Data and code used in this manuscript can be accessed here: https://github.com/goldrieve/vsgseq2.

### Software availability

The vsgseq2 pipeline can be accessed here: https://github.com/goldrieve/vsgseq2. The version described in this manuscript is v.1.0.2

## Competing interests

No competing interests were disclosed.

## Grant information

Wellcome Trust (221717/Z/20/Z) Wellcome Trust (206815/Z/17/Z)

## Acknowledgements

We thank Dr Alex Beaver, Dr Jane Munday, Prof. Liam Morrison and Prof. Richard McCulloch for their valuable insights on this manuscript.

## Figures and Tables

**Table 1:** Updates in tools used by vsgseq2 broken down by each step in the pipeline (Fig. 1).

| Dependency | VSGSeq | vsgseq2 |
| --- | --- | --- |
| Repository | https://github.com/mugnierlab/VSGSeqPipeline | https://github.com/goldrieve/vsgseq2 |
| Language | Python (v2.7.18) | Python (v3.10.11), R (v4.3.2), Nextflow (v.25.04.1.5946) <sup>10</sup> |
| TRIM | trim_galore (v.0.6.10) <sup>14</sup> | trimmomatic (v.0.39) <sup>15</sup> |
| ASSEMBLE | Trinity (v2.8.5) <sup>16</sup> | Trinity (v2.15.1) <sup>16</sup> |
| ORF | VSGSeq <sup>4</sup> | TransDecoder (v.5.7.1) <sup>17</sup> and seqkit (v.2.8.0) <sup>18</sup> |
| BLAST | blastn (v.2.7.1) <sup>19</sup> | blastn (v.2.16.0) <sup>19</sup> |
| CLUSTER | cd-hit-est (v.4.8.1) <sup>20</sup> | cd-hit-est (v.4.8.1) <sup>20</sup> |
| INDEX / MAP | bowtie (v.1.2.3) <sup>21</sup> | Salmon (v.1.10.2) <sup>11</sup> |
| QUANTIFY | MULTo (v.1.0) <sup>22</sup> | Salmon (v.1.10.2) <sup>11</sup> |
| SUMMARISE | VSGseq <sup>4</sup> | vsgseq2 and multiqc (v1.21) <sup>23</sup> |

## Notes

https://github.com/goldrieve/vsgseq2

### Competing Interest Statement

The authors have declared no competing interest.

